# TopoDiff: Improving Protein Backbone Generation with Topology-aware Latent Encoding

**DOI:** 10.1101/2023.12.13.571602

**Authors:** Yuyang Zhang, Zinnia Ma, Haipeng Gong

## Abstract

The *de novo* design of protein structures is an intriguing research topic in the field of protein engineering. Recent breakthroughs in diffusion-based generative models have demonstrated substantial promise in tackling this task, notably in the generation of diverse and realistic protein structures. While existing models predominantly focus on unconditional generation or fine-grained conditioning at the residue level, the holistic, top-down approaches to control the overall topological arrangements are still insufficiently explored. In response, we introduce TopoDiff, a diffusion-based framework augmented by a global-structure encoding module, which is capable of unsupervisedly learning a compact latent representation of natural protein topologies with interpretable characteristics and simultaneously harnessing this learned information for controllable protein structure generation. We also propose a novel metric specifically designed to assess the coverage of sampled proteins with respect to the natural protein space. In comparative analyses with existing models, our generative model not only demonstrates comparable performance on established metrics but also exhibits better coverage across the recognized topology landscape. In summary, TopoDiff emerges as a novel solution towards enhancing the controllability and comprehensiveness of *de novo* protein structure generation, presenting new possibilities for innovative applications in protein engineering and beyond.

## 1. Introduction

The *de novo* design of proteins refers to the task of designing a physically plausible protein backbone and the corresponding sequence without involving naturally occurring proteins as starting points [1], which is an intriguing field of research with the potential to venture into unknown structure space, offering limitless opportunities for tailoring proteins to novel applications [2]–[7]. Despite its vast potential, the *de novo* protein design has long been recognized as a challenging task, considering the highly-structured nature of proteins and the stringent requirements on physical restraints [8], [9].

People have used Generative Adversarial Networks (GAN [10]) [11] and Variational Autoencoders (VAE [12]) [13]–[17] to model the high-dimensional structural space. Those models typically represent the proteins as 2D distance matrices and the outputs were often a predicted distance matrix that frequently suffered from inconsistency issues [18]. Despite the limitation, VAE-based models showed promise in learning a valuable latent space to reduce the dimensionality of the problem, while also offering increased controllability and interpretability, such as incorporating a differentiable coordinate-based heuristic with VAE’s generative prior for constrained structure generation [16] and modifying the protein structure via interpretable manipulations over the latent code [15].

Recent advances in the diffusion models significantly reshaped the field with its superior ability to generate novel, diverse and physically plausible structures. Though the pioneer efforts still relied on 1D or 2D protein representations [19]–[21], subsequent works tended to build an equivariant network to directly learn the physical prior in the Cartesian space [22]–[28]. Despite the encouraging results, previous methods primarily focused on generating structures unconditionally or only incorporated localized conditioning at the residue level [22], [26]. To the best of our knowledge, the only attempt on global-structure conditioning [27] employed a mechanism methodologically resembling the classifier guidance [29], which requires the use of well-trained external classifiers, and fundamentally, the availability of appropriate labels for classifier training. Hence, it remains insufficiently explored how to learn a meaningful latent space for the global structures of proteins and to enforce global controllability correspondingly during the structure generation. Furthermore, while the potential issue of mode collapse in some diffusion-based models has been noticed [25], the current metrics provide no indication of the extent to which the natural protein space has been covered.

In this work, we seek to bridge the gap between VAE-based models and diffusion-based models, leveraging the strengths inherent in both approaches. Our contributions are as follows:

- We propose a fully equivariant encoder-decoder network to jointly learn a global structure-level latent space and a generative module conditioning on such information.
- We adapt a coverage metric to quantitatively measure the coverage of sampled structures over the natural protein fold space and demonstrate its effectiveness in addressing underrepresented topologies.
- We show that the acquired latent space is highly structured and interpretable, and that the generative module conditioning on it could achieve enhanced coverage of natural topologies and comparable performance on other established metrics.
- We demonstrate some primary usages of the latent space towards controllable structure generation.

## 2. Methods

### Diffusion model

With *R* denoting the protein with length *l*, we define the forward process of the diffusion model as a non-learnable Markovian process that gradually introduces noise to a protein structure ℛ ^0^ towards a pre-defined prior distribution *p*_*T*_ . The reverse process is also a Markovian process that learns to remove the noise signal, with a fixed-sized latent variable **z** as a globally aware condition, representing the structure latent of the ground truth data ℛ ^0^.

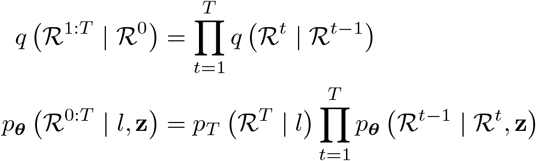

The training objective (Figure 1a) is to predict the ground truth ℛ ^0^, which is, for simplification, denoted here as reconstruction loss 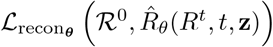. The detailed formulation of the forward and reverse transition kernels *q, p*_***θ***_, and the loss 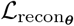 are provided in Appendix A.1.

**Figure 1:**
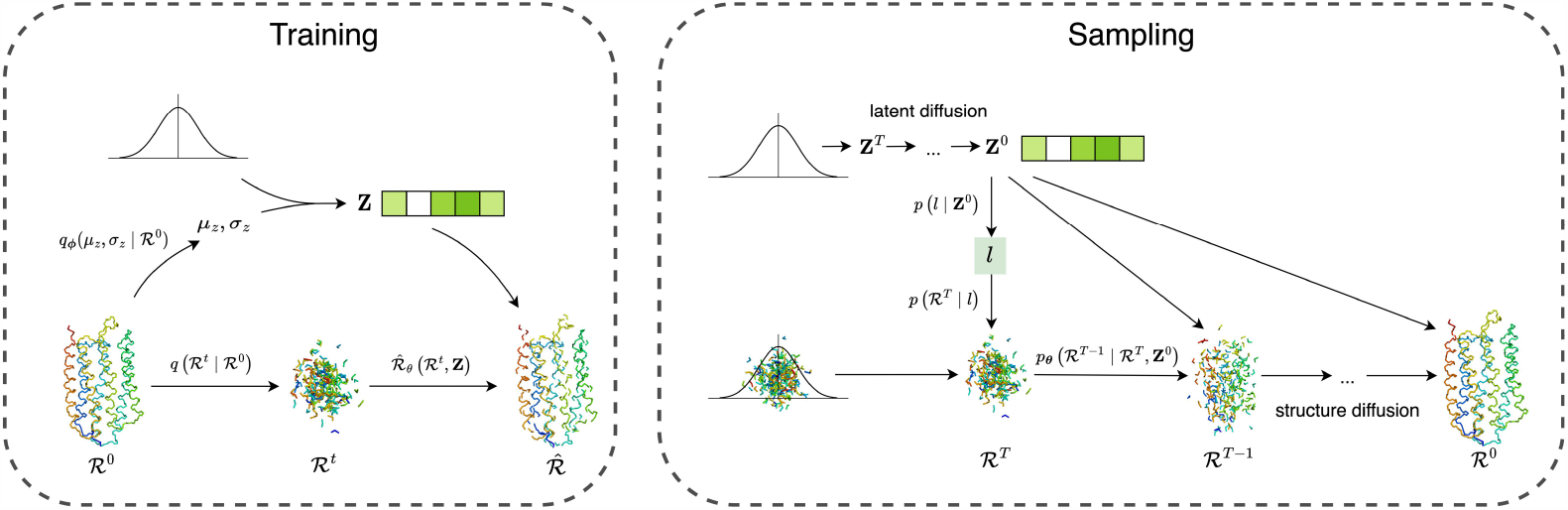
Illustration of the training and sampling process with TopoDiff

### Topology encoder and the VAE training objective

We incorporate an additional encoder module designed to learn the fixed-size latent topology representation **z** for a given data ℛ ^0^. We model the *p*(**z**) with an isotropical Gaussian prior, and add a KL divergence loss term to encourage a continuous latent structure, effectively shaping our model into a VAE-like framework. Combined with the diffusion model, our final training objective is:

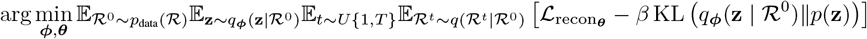

Additional details of the encoder implementation are provided in Appendix A.2, and the derivation of the evidence lower bound (ELBO) justifying this training objective is provided in Appendix A.6.

### Latent diffusion model over the learned latent space

Motivated by previous works [30], [31], we train an additional latent diffusion network to model the latent distribution. Further implementation details are available in Appendix A.3.

### Sampling process

We briefly summarize the process of sampling with the model. For unconditional sampling (Figure 1b), we first sample **z** with the latent diffusion model. Then, we sample the protein length *l* with a straightforward KNN regression scheme, by querying the *k* nearest neighbours in the latent space from the training dataset: 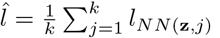 and *l ∼* Uniform 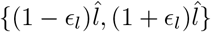 where the hyperparameter *ϵ*_*l*_ is engaged to decide the range of variation. Finally, we use (**z**, *l*) as the condition to conduct the structure sampling described in Methods 2. For fixed-length sampling, we resample latent until the target length falls within the range of variation.

### Metrics for sample quality evaluation

We employ a variety of metrics to comprehensively evaluate the performance of a model during unconditional sampling. Here, we briefly describe the choice for the newly proposed coverage metric. The exact definitions of all metrics are described in Appendix A.4. To measure the extent to which our model can cover the natural protein space, we adapt the coverage metric initially proposed by Naeem et al. [32]. Briefly, we first construct a KNN manifold of all real samples (*i*.*e*. natural proteins) and then measure the fraction of real samples whose neighbourhoods encompass at least one fake sample (*i*.*e*. structures produced by the generative models). Formally, with *{ X*_*i*_ *}*denoting the set of real samples, *{ Y*_*i*_ *}* denoting the all fake samples, NND_*k*_ (*X*_*i*_) denoting the distance from *X*_*i*_ to its *k*^*th*^ nearest neighbour in *{ X*_*i*_ *}*excluding itself, and *B*(*x, r*) denoting the hypersphere around *x* with a radius of *r*, the coverage is defined as:

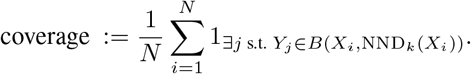

## 3. Experiments

### 3.1 Interpretability of the global-structure-centric latent space

To gain an understanding of the latent space learned by the model, we encode all structures in the CATH-60 [33] dataset into the 32-dimensional latent codes and visualize with t-SNE. As shown in Figure 2, the latent codes altogether form a compact and continuous manifold. Interestingly, the clustered structures perfectly coincide with the human curation of CATH class annotations, with each class clearly separable from the others in the 2-dimensional space. Besides, although the latent space does not necessarily correspond perfectly with human annotation at the finer level, we find that each CATH Architecture cluster indeed exhibits a unique spatial distribution (Figure 4). These observations indicate that the latent space obtained in this work is highly interpretable. The model learns to partition over the structure space in an unsupervised manner, forming a continuous while structured manifold that encodes the essential information about the underlying protein topologies, which may serve as informative conditions to the downstream generative module for controllable structure generation.

**Figure 2:**
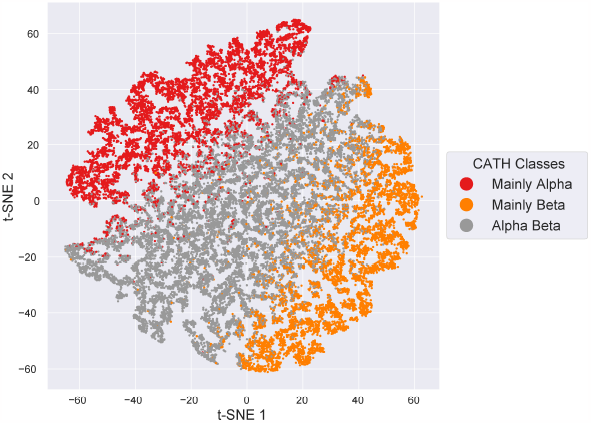
t-SNE visualization of the latent space

### 3.2 Results on Unconditional Generation of Monomer Proteins

We further conduct a comprehensive evaluation on the performance of our generative module for unconditional protein structure sampling against other state-of-the-art models. For each model, we randomly generated 500 structure samples at each fixed length of 50, 75, …, 225, 250, a series uniformly spanning the range of our training data, and measured the performance respectively.

As depicted in Figure 3, TopoDiff exhibits some unique characteristics and advantages. In terms of the coverage of natural protein topologies, TopoDiff ranks the top among all sampled lengths, showing the effectiveness of modeling the natural data distribution with a structure-centric latent space. To further investigate the exact advantage of our model in mode coverage, we analyze the sample-wise binary coverage indicators and find that our model could encompass a significantly larger number of natural folds with mainly beta-strand compositions, a group of topologies that are typically underrepresented by other methods (Appendix C, D). Regarding designability, TopoDiff is comparable to other models with similar parameter sizes (Genie [25] and FrameDiff [24]), and gradually exhibits advantages as the length increases. As for novelty, TopoDiff maintains a fair balance between the coverage of known topologies and the generalizability to novel topologies, with the median value of the maxTM consistently around 0.6. In Figure 10, we present several randomly selected novel structures with considerable confidence in designability.

**Figure 3:**
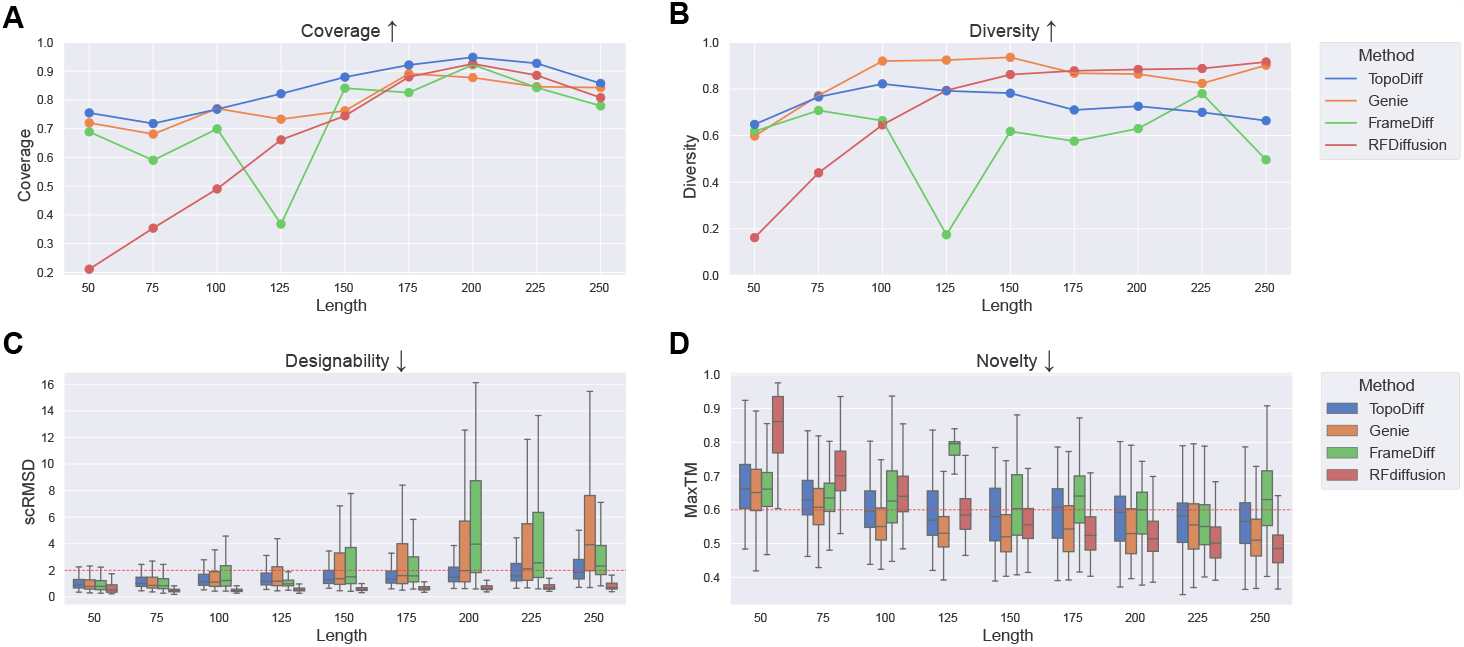
Evaluation of the models for unconditional sampling

**Figure 4:**
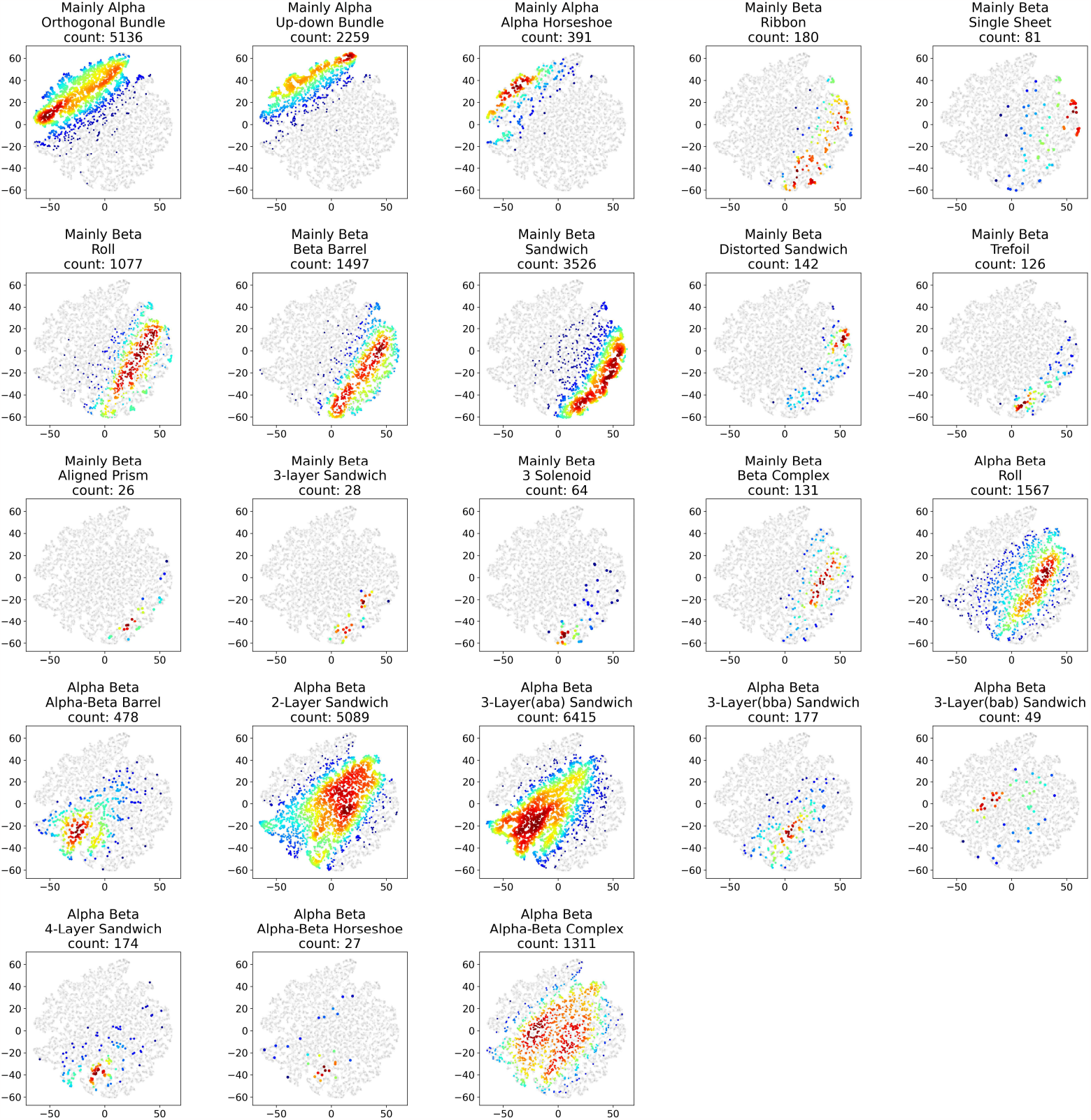
Density plot of each CATH architecture in the t-SNE dimension-reduced latent space

**Figure 5:**
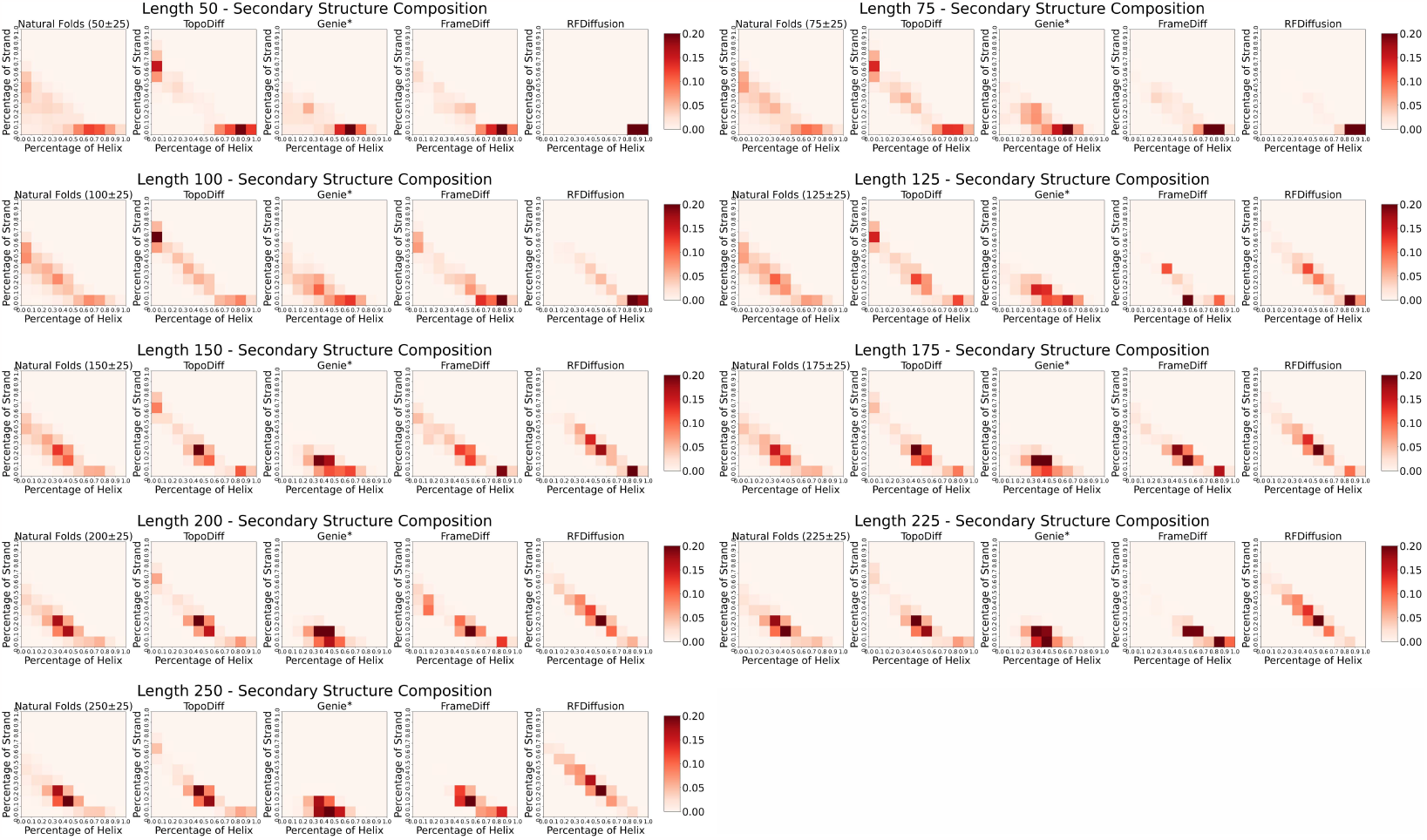
Comparison of the distribution of secondary structure composition at each length range. For each sampled length, we calculate the secondary structure composition for each sample and visualize the distribution of both natural proteins and the four models. The samples generated with our model consistently exhibit strong alignment with the natural distribution.

### 3.3 controllable Generation with the Latent Encoding

Since the topologies of sampled structures are primarily determined by the latent encoding in our method, we could achieve the controllable generation of the protein structures by simply manipulating the latent code fed to the decoder. Importantly, since the latent code is sampled from a lowdimensional compact distribution, the time for such manipulations is almost negligible. Hence, in TopoDiff, controls could be efficiently exerted over the protein topology, prior to exhaustive sampling at the residue level in the Cartesian space.

#### Controlling the Trade-Off Between Sample Designability and Coverage/Diversity

We find that the latent encodings are not equally likely to be decoded into designable proteins. Interestingly, the likelihood exhibits a specific pattern in the latent space. Based on the scRMSD measurements of the unconditional sampling result in the first run, we train a prediction layer to forecast the expected outcome of a given latent, and then conduct rejection sampling with the prediction layer with different cutoffs. With this sampling scheme, we could effectively tune the trade-off between designability and coverage/diversity, and prohibit the model from sampling those highly disordered topologies that comprise a certain proportion of the training set. The detailed experiment design and result are provided in Appendix F.

#### Sampling Similar Topologies With Respect to A Given Query Protein

The compact and continuous nature of the latent space enables the generation of a plethora of artificial sample structures around an arbitrary query protein. To test this capability, we select representative proteins of different annotated architectures from the training dataset, calculate their latent encodings, and then randomly sample the latent space using the inferred posterior distribution. As shown in Appendix Figure 12, with a temperature of 0.1, the artificial samples produced by TopoDiff exhibit similar spatial arrangements of secondary structures, and considerable residue-level diversity with the query proteins. The result demonstrates that TopoDiff, with its explicitly modeled latent topology space, could provide additional global controllability for structure sampling.

## 4 Discussion

In this work, we propose a general framework for simultaneous learning of a latent representation for the protein global structures and training of a generative module for atomic-level backbone generation, all within a completely unsupervised context. The inherent equivariance of the encoder and decoder endows the model with the capacity to learn and reconstruct crucial structural variations directly in the Cartesian space, thus circumventing the potential information loss or bias when converting to other structural parameterizations such as the inter-residue distances. The learned fixed-size representation encodes essential information of the global structures, altogether forming a compact and sequence-agnoistic latent space, which holds the potential to advance our comprehension of the protein structure universe and lays the foundation for controllable structure generation at the same time. With the help of the latent diffusion module, we could achieve unconditional sampling over the whole space and discover novel topologies with ease.

We also introduce a novel metric to quantitatively measure the coverage of sampled structures over the established protein structure space. The ability to cover all known topologies is a useful indicator to ensure that the model does not suffer from significant mode collapse, which has been observed in some models particularly within mainly-beta categories but still lacks thorough analysis [25]. Furthermore, as demonstrated in Appendix D, the metric could be used not only as a global indicator, but also in a sample-wise manner to pinpoint underrepresented regions, showing promise in quantifying the sampling limitations of the existing models and guiding future improvements. While we currently use a third-party model [34] to measure the pair-wise distance between structures for the implementation of the above metric, it is worth noting that other distance functions can be similarly utilized, offering flexibility in its implementation. We anticipate that the development of this metric could facilitate the systematic evaluation of generative models in protein design.

A distinctive highlight of our work is the latent encoding learned by the model, which can serve as a control over the overall topology. This control paradigm stands orthogonal to the residue-level finer-grained control [11], [26]. While finer-grained control holds utility in numerous situations, the manual assignment of residue-wise condition demands domain expertise and might limit the exploration space of sampling. The global control, in contrast, resembling a top-down approach to control the designed topology, offers advantages in scenarios where both topology-level control and residue-level variation are desirable [35], [36]. We believe that the combined utilization of global and residue-wise conditions could represent a potent approach for achieving controllable *de novo* design.

## Acknowledgement

We would like to thank Jian Hu and Zefeng Zhu for helpful discussions during the course of this work.

## Appendix

### A. Implementation detail

#### A.1. Implementation detail of the diffusion model

##### Parameterization of the protein

In alignment with prior research [22]–[26], we represent the protein as residue clouds in the SE(3)^*N*^ space. Briefly, for a sequence of length *l*, each residue is parameterized as the collection of the translation of its C_*α*_ atom, denoted as **t**_*i*_ *∈* ℝ^3^, and its orientation, uniquely defined by the coordinate of three atoms denoted as **r**_*i*_ *∈* SO(3). Collectively, we denote the whole sequence as *R* = *{*(**t**_*i*_, **r**_*i*_)*} ∈* SE(3)^*l*^.

##### Diffusion on ℝ^3^ space

For translation coordinates **t**_*i*_, the diffusion process closely follows the original setup of DDPM [37], where the coordinates are gradually perturbed towards *𝒩* (**t**_*T*_ ; ***O*, I**).

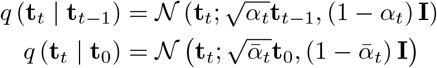

The model is trained to predict the ground truth coordinates at *t* = 0, and the reverse process is given by:

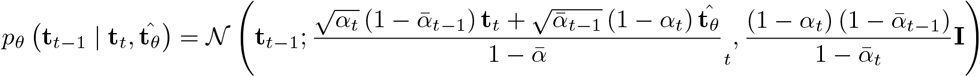

##### Diffusion on SO(3) space

For SO(3) space, the forward transition is formulated to compose a perturbation with the ground truth orientation, which conforms to ℐ𝒢_*SO*(3)_ distribution. Practically, we represent the perturbation with the axis-angle representation (*ω*, **v**), where *ω* is sampled from the marginal distribution of *ℐ 𝒢*_*SO*(3)_ in radians, and **v** is uniformly sampled from *S*^2^ unit sphere.

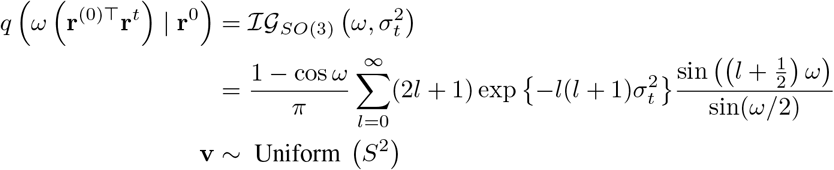

Following [23], we empirically formulate the reverse transition in an iterative perturb-denoise scheme.

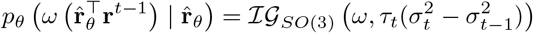

*{ σ*_*t*_*}, { τ*_*t*_*}* are two fixed schedules that determine the noise scales for each transition.

##### Model architecture

The model backbone we use is adapted from [24], which comprises an embedder module for all conditional information (*e*.*g*., timestep, residue index, topology latent) and a structure module with 4 IPA layers for structure prediction. The total timestep is set to 200.

##### Training objective

The training objective is to predict the ground truth *ℛ*^0^ out of the noised data *ℛ*^*t*^.

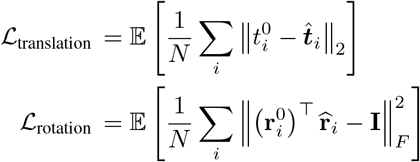

In addition, we use fape loss ℒ;_fape_ and distogram loss ℒ_distogram_ as auxiliary losses to stabilize the training, which is defined the same as [38].

#### A.2. Implementation detail of the topology encoder

Inspired by [34], we use six-layer EGNN [39] to extract the local topology information and incorporate two layers of transformer encoder for information integration and pooling.

#### A.3. Implementation detail of the latent diffusion model

The formulation of the latent diffusion mirrors the translation diffusion on ℝ^*N*^ space. We use a backbone of a 10-layer MLP with skip-connections as the model backbone. The total time step is set to 200.

#### A.4. Implementation detail of the evaluation metrics

##### Designability

To assess the designability of a given sample, we first use ProteinMPNN [40] to sample 8 amino acid sequences with a temperature of 0.1. Subsequently, the sequences are fed to ESMFold [41] to infer the structures. The minimum RMSD of the inferred structures to the given sample is reported as scRMSD.

##### Novelty

To assess the novelty of a given sample. We begin by using Foldseek [42] to query the sample against the CATH-40 dataset [33] with the parameter “-a 1 –exhaustive-search 1 -e inf –c 0.5 –alignment-type 1”. As Foldseek employs a slightly different implementation of TM-align [43], we subsequently select the top 25 matches from the query results with the highest TM-scores and re-compute the alignment with TM-align [43]. The highest TM-score to the chains in the dataset is reported as maxTM, representing the novelty of a sample (a higher score implies the sample is generally less novel).

##### Diversity

To compute the diversity of *N* samples, we first use TM-align [43] to compute the pairwise TM-scores. Then, we cluster the samples with a cutoff of 0.6. The proportion of total clusters to the total number of samples *N* is reported as diversity.

##### Coverage

As is described in Methods 2, for *N* given samples, the coverage over target sample distribution is defined as:

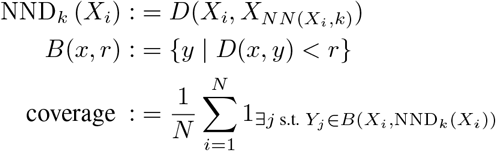

where *{ X*_*i*_ *}* denotes the set of real samples, *{ Y*_*i*_ *}* denotes the set of all fake samples, NND_*k*_ (*X*_*i*_) denotes the distance from *X*_*i*_ to its *k*^*th*^ nearest neighbour in *{ X*_*i*_*}*excluding itself, and *B*(*x, r*) denotes the hypersphere around *x* with a radius of *r*.

Based on this definition, we need to define a function *D*(·, ·) to measure the distance of two arbitrary structures. Specifically, we need to compute at least the distance between *X*_*i*_ and the *k*^*th*^ nearestneighbours in *{X*_*j*_ | *j ≠i}* to construct the KNN manifold at the given point, and then use the distance between *X*_*i*_ and its 1^*st*^ nearest neighbour in *{Y*_*i*_*}* to decide if *X*_*i*_ is covered by an arbitrary fake sample. The final coverage metric is obtained by averaging the binary indicators over *{X*_*i*_*}*.

As *k* increases, the accurate calculation of such distances can be exceedingly time-consuming when using traditional structural alignment algorithms (*e*.*g*., TM-align), which involve a series of dynamics programming and heuristic iterative algorithms to refine optimal solutions. Therefore, we resort to a third-party tool [34], which employs supervised contrastive learning to learn a metric for structure comparison. In practice, we read the coordinates of all samples, encode them with the model to get their vectorized representations, and then compute the pairwise cosine similarities, following the designated usage of the model.

The choice of *k* is a hyperparameter that should be determined prior to the computation of the metric. Following the recommendation of the original paper [32], we choose *k* such that a sample size equivalent to the artificial samples from the very same distribution of real samples could achieve a coverage close enough to 1. To do this, we initially randomly sample 500 natural chains as the pseudo-query set, use the remaining chains as samples from the target distribution, and then compute the coverage of the pseudo-query set against the target distributions for different choices of *k*. Based on such a scheme, we ultimately use *k* = 100 for all experiments, although we find different choices of *k* generally do not alter the relative rankings of the evaluated models.

When comparing the samples of a fixed length to the natural protein distribution, at each sampling length *l*, we consider all natural protein structures in the CATH-40 dataset [33] lying within the interval of [*l −*25, *l* + 25].

#### A.5 Training

##### Dataset

We use the CATH-60 non-redundant annotation list [33] for the training of the model. For each sample (chain), we first trim the leading and tailing residues that lack ground truth coordinates to get the effective length of a sample. We then subset the dataset to contain only samples within the range of [50, 256]. We also remove samples with more than 20% inner gaps or over 60% disordered regions. Following these data preprocessing steps, our training dataset comprises a total of 30,074 samples.

#### A.6 Derivation of the evidence lower bound (ELBO)

##### Modeling the protein space with variable length

The training dataset frequently encompasses protein samples of varying length. To standardize the representation of these structures, we denote the maximum residue length in the dataset as *L*_max_. For a training sample with length *l*, we could uniquely map it to the higher dimensional manifold 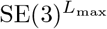, with the first *l* frames representing its coordinate at SE(3)^*l*^ manifold, followed by *L*_max_ *− l* padding frames.

Under this parameterization scheme, the prior distribution *p*_*T*_ (ℛ ^*T*^ | *l*) characterizes a process that perturbs the first *l* frames in accordance with the respective prior distribution, leaving the rest of frames padded. Thus, for any timestep *t, q* (*l*| ℛ ^*t*^) is the delta distribution concentrated at 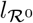, as the length is naturally determined by the number of non-padded dimensions.

##### Modeling the joint distribution of data and intermediate variables

In accordance with the sampling process outlined in Figure 1, we could now formulate the joint distribution of all variables involved in structure sampling as below:

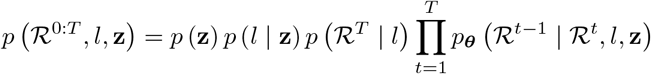

##### Evidence lower bound

We could eventually derive the ELBO of our diffusion-VAE model.

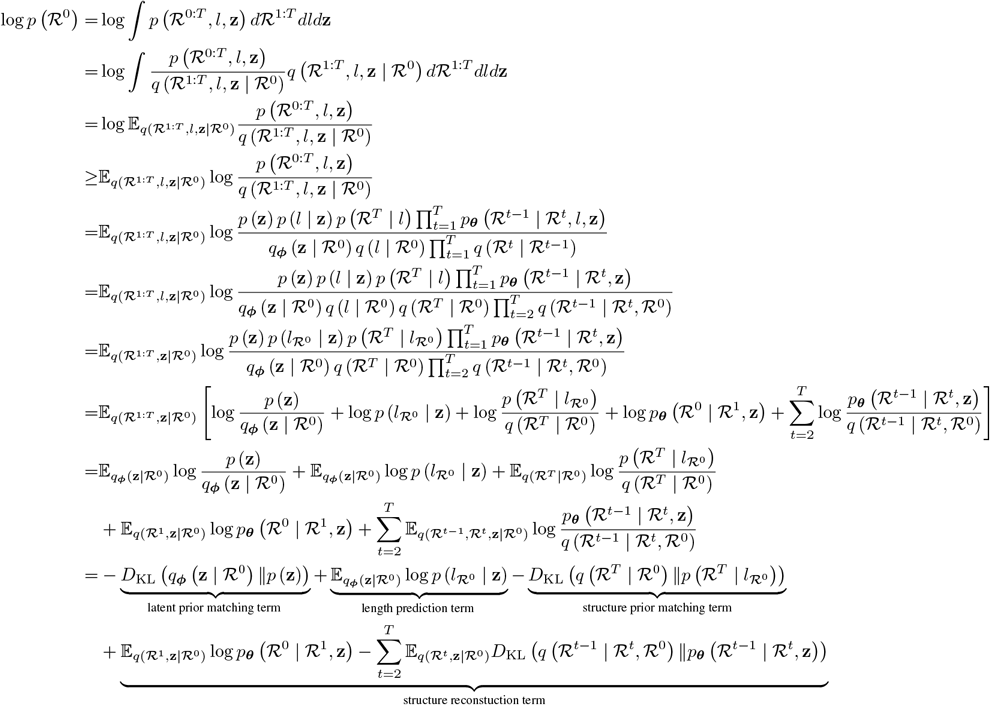

The structure prior matching term has no trainable parameters and is fulfilled naturally with our pre-defined noise schedule. In the current implementation, we estimate the length distribution with a KNN approach described in Methods 2, which is empirically sufficient and also non-learnable. Therefore, our training objective is to minimize the collection of latent prior matching term (the KL divergence between the posterior and prior of **z**) and the structure reconstruction term.

### B Visualization of the latent space with CATH Architecture annotations

### C Comparison of the distribution of secondary structure composition

### D Using coverage metric to find underrepresented natural folds in generated samples

When computing the coverage, besides the final scalar bounded between 0 and 1 indicating the extent to which the target distribution has been covered, we could also examine the sample-wise binary indicator to gain a deeper understanding of which specific modes are covered and which are not. Here, we illustrate this analysis using the artificial samples with the chain length of 125.

First, we present a Venn diagram for the four models under evaluation to visualize the overlapping coverage of natural folds (Figure 6). Out of the 9515 natural folds within the range of [100, 150], 2659 folds are commonly covered by all of the 4 models (region *a*), and a total of 5871 folds are covered by at least 3 models. Our model stands out by having a unique coverage over 757 folds (region *b*), a number even remarkably higher than the sum of all folds uniquely covered by the other three models. Additionally, there are also 1074 natural folds that are not considered as covered by any of the four models (region *c*).

**Figure 6:**
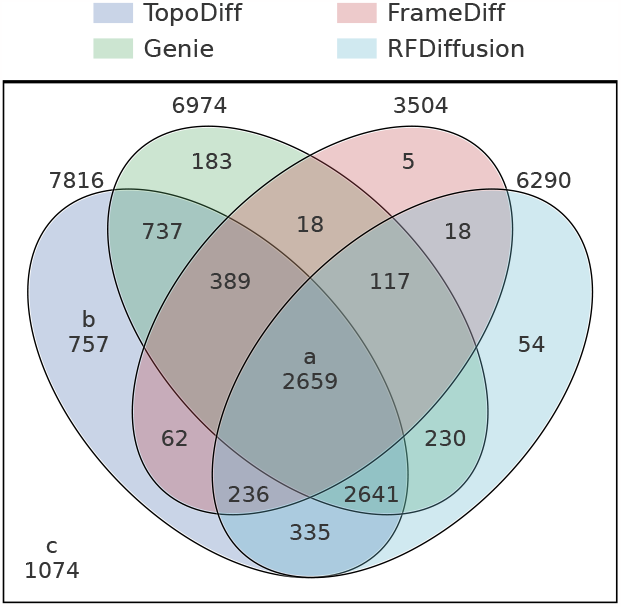
Venn diagram of covered natural folds with different methods (length 125)

To gain a more intuitive understanding of the characteristics of the regions of samples stated above (*e*.*g*., region *a, b, c*), we randomly select 50 samples from each region and provide visualization as below. For the commonly covered natural folds (region *a*, Figure 7), most of them consist of a high proportion of alpha helices and are generally well-packed into a globular shape. Further examination of the samples uniquely covered by TopoDiff (region *b*, Figure 8) reveals that our model could cover a great number of natural folds characterized by a mainly beta composition. Moreover, most of the commonly uncovered folds (region *c*, Figure 9) are still proteins of this category, with some adopting even more complex global arrangements. There are also some uncovered highly stretched up-down bundles, which are unlikely to be useful in real design scenarios. Overall, our result demonstrates that there is still room for improvement in the current generative models on the successful modeling of complex and diverse proteins with the mainly beta composition.

**Figure 7:**
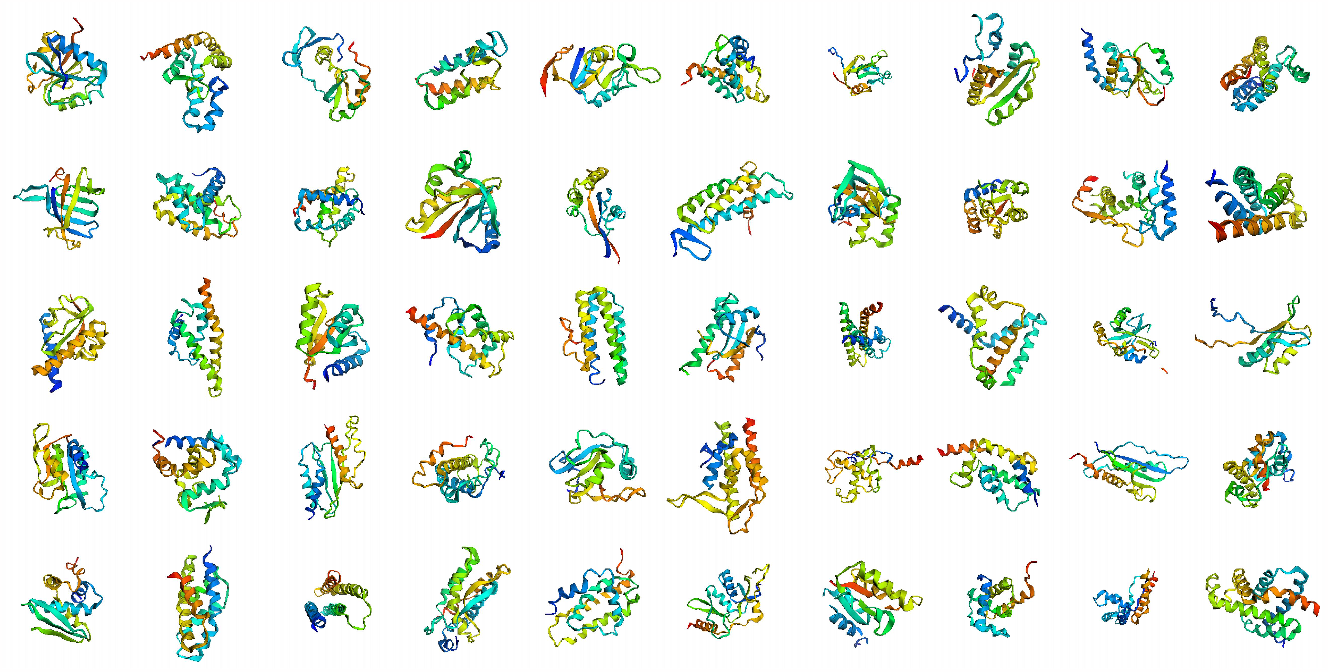
Examples of commonly covered natural folds

**Figure 8:**
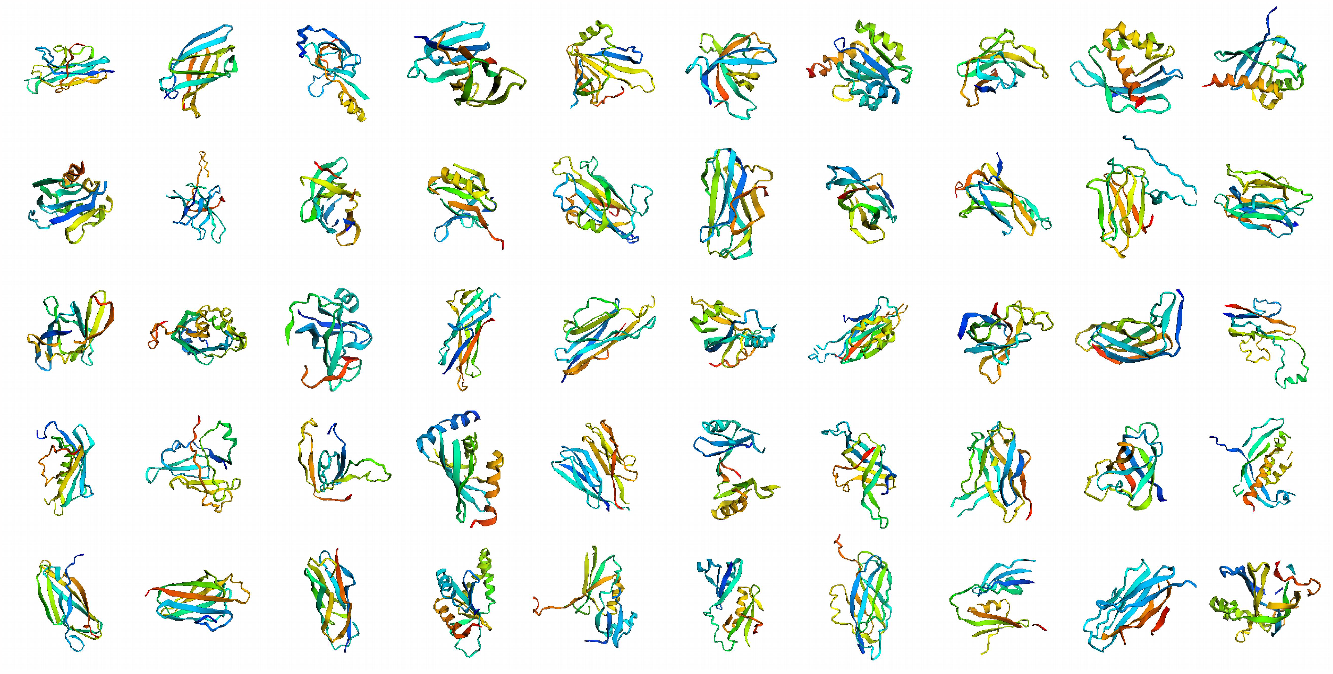
Examples of uniquely covered natural folds by TopoDiff

**Figure 9:**
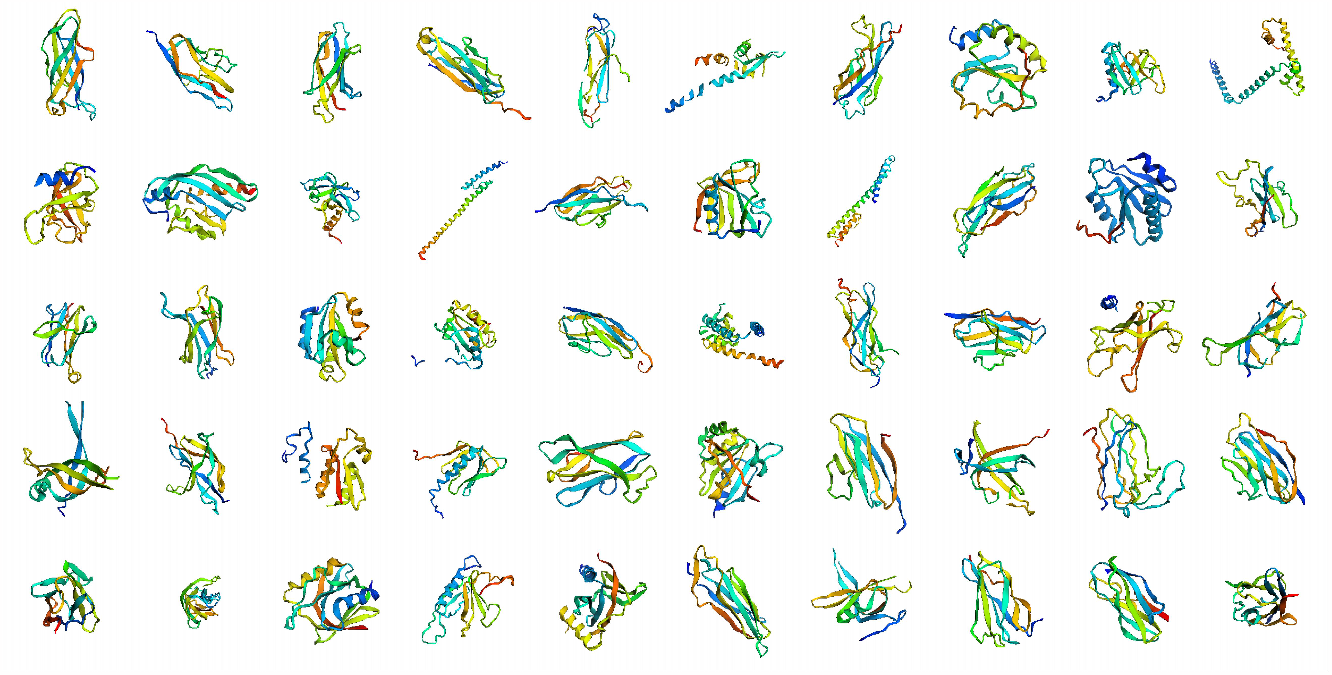
Examples of commonly uncovered natural folds

**Figure 10:**
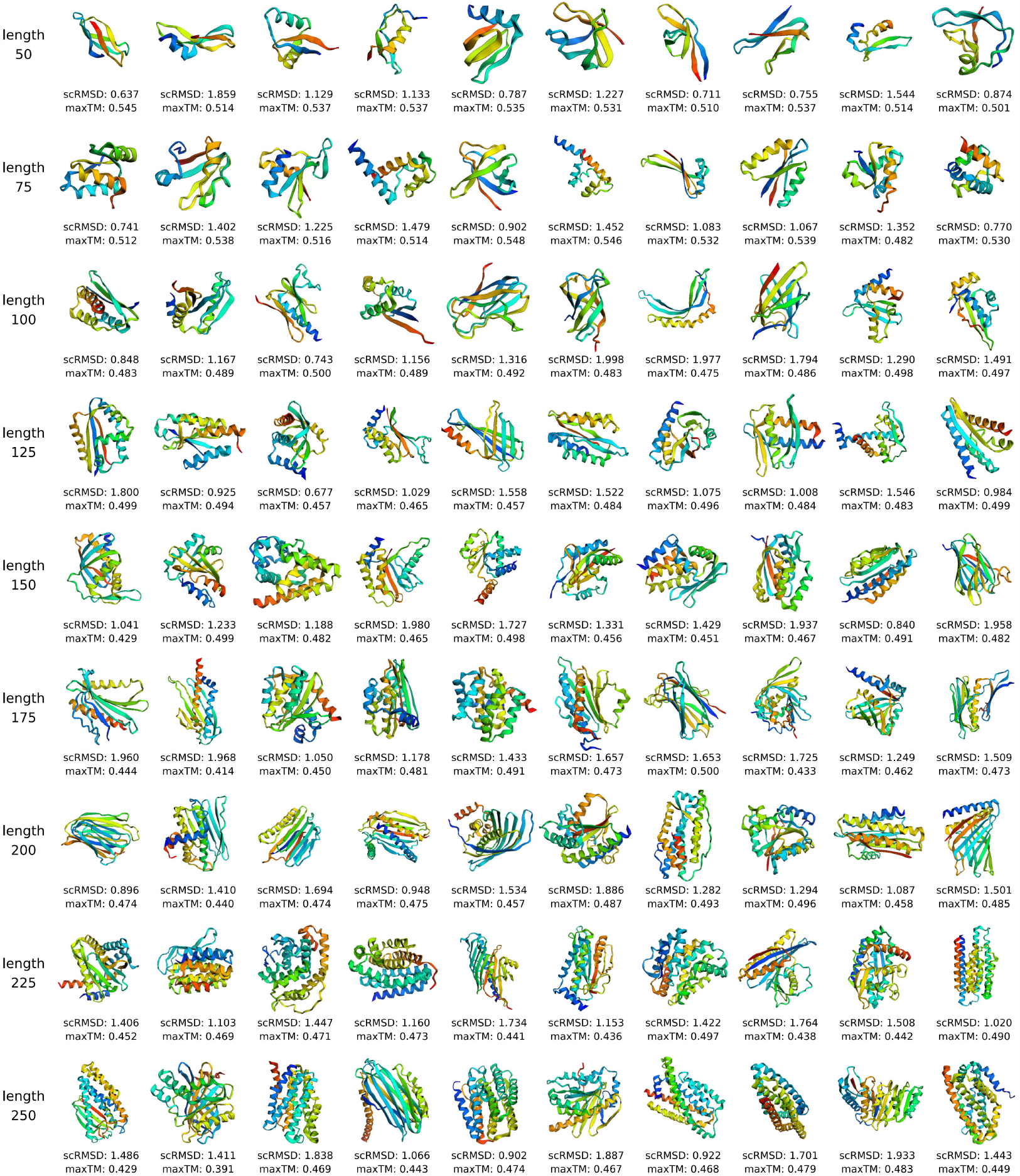
Randomly selected novel folds with confidence in designability. Using maxTM *<* 0.5 (maxTM *<* 0.55 for length 50 and 75) and scRMSD *<* 2Å as cutoff, we subsample a series of novel folds with confidence in their designability. Here, we randomly select 10 samples per length for visualization.

This demonstration highlights the utility of the coverage metric in identifying uncovered folds and detecting potential regions of mode collapse, which may provide valuable insights for model improvement.

### E Examples of novel proteins in generated samples

### F Controlling the trade-off of designability and coverage/diversity

Specifically, for each chain length in the range of [50, 250], we randomly sample 10 structures, ensuring broad coverage across the entire latent manifold, and measure the scRMSD values of these samples. We then collect all measurements to train a simple prediction network that forecasts the expected scRMSD for a given latent encoding, and finally use this prediction as a cutoff to refrain from sampling in the region of latent space with predicted scRMSD higher than this cutoff.

As shown in Figure 11, by adjusting the cutoff over predicted scRMSD, we obtain various models with different performance trade-offs between designability and diversity/novelty/coverage. Therefore, instead of the traditional computationally intensive approach of exhaustive sampling directly at the residue level in the Cartesian space followed by post-hoc selection, our method allows the learning of the relationship between the latent encodings and expected outcomes, through which conditional sampling could be performed in the latent space to finely tune the trade-off between designability and diversity/coverage *a priori*, a scheme that is significantly less costly in computation considering the low-dimensionality of the continuous latent space.

**Figure 11:**
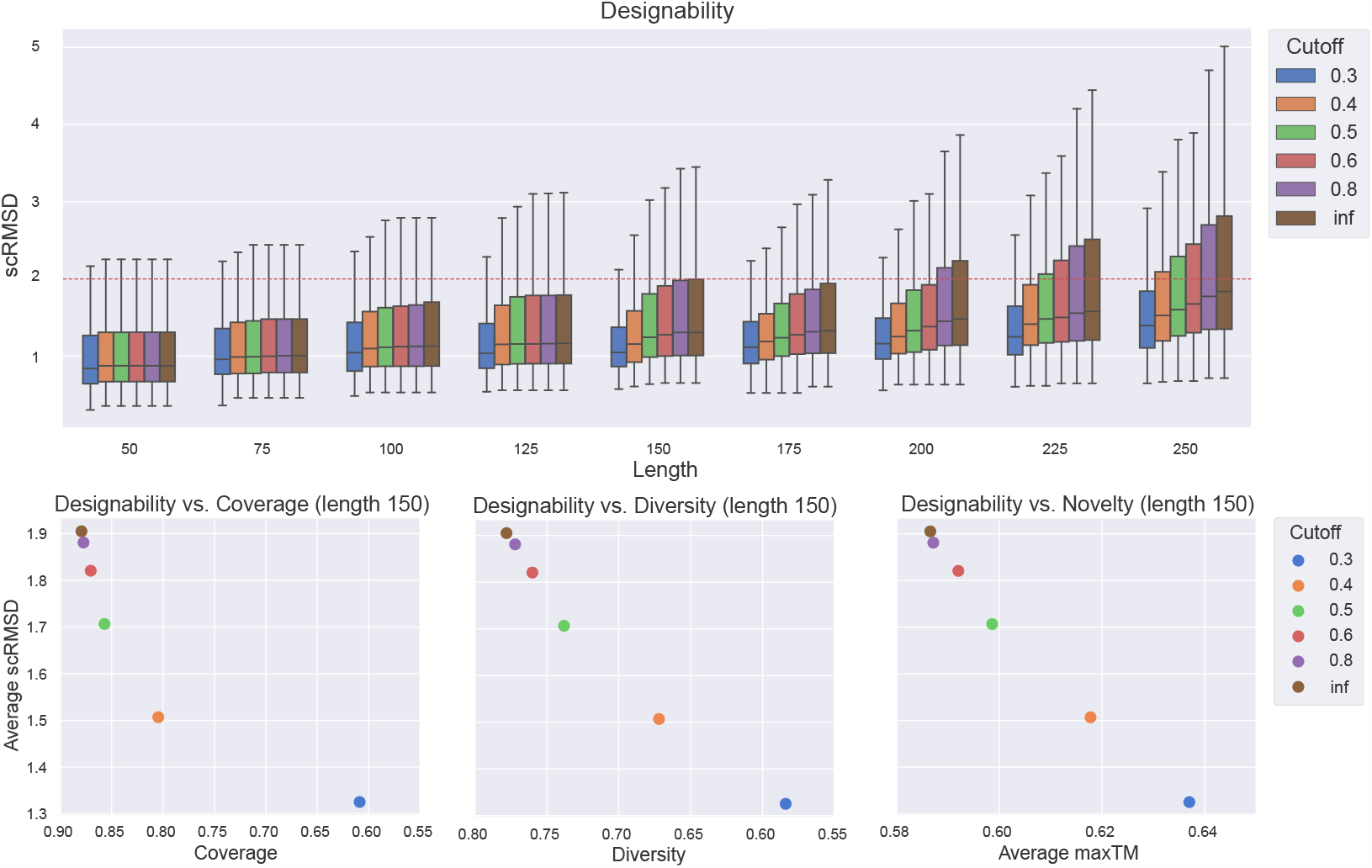
Controlling the trade-off between sample designability and coverage with latent space rejection sampling

**Figure 12:**
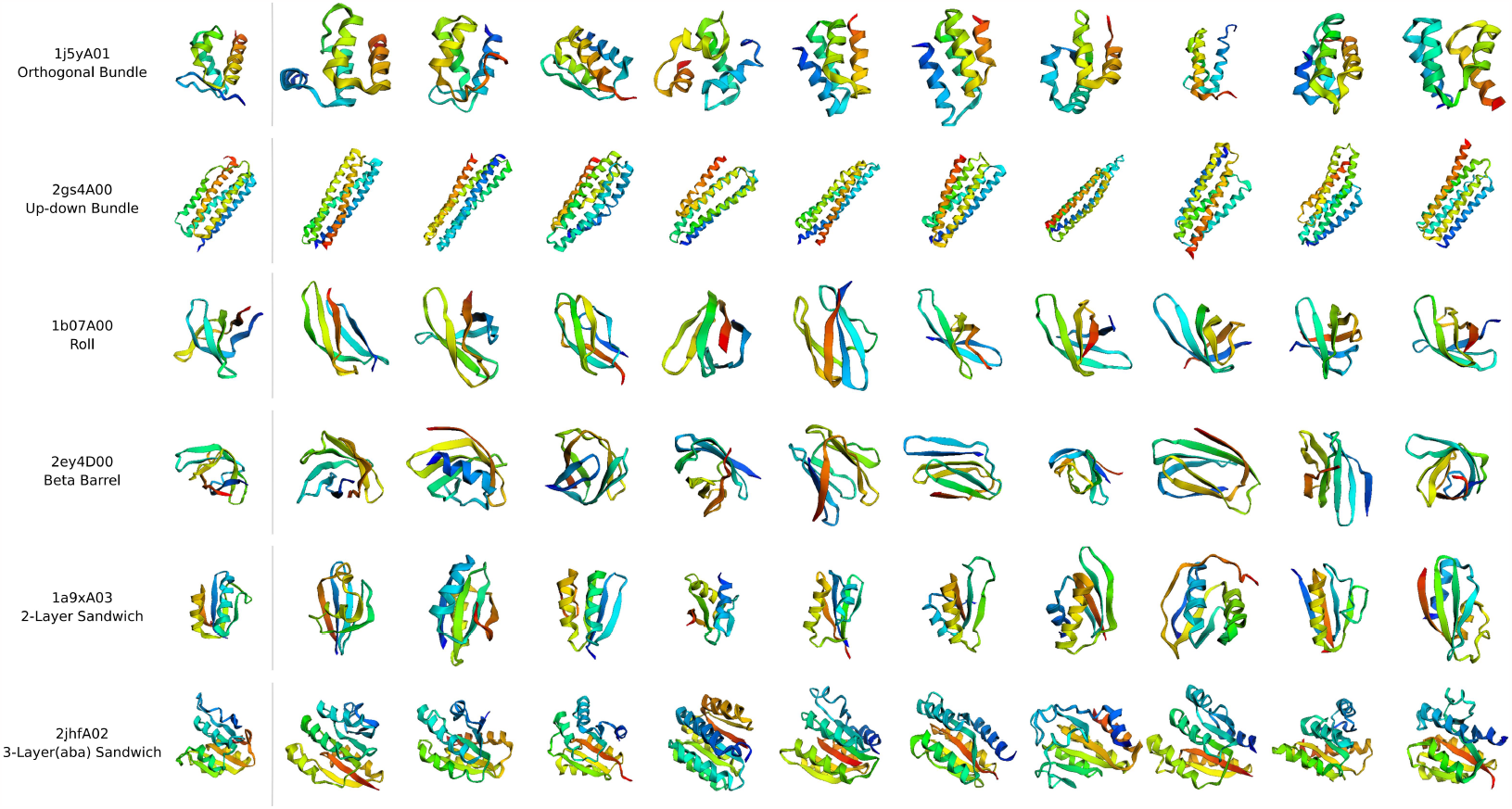
Sampling proteins with similar topology with respect to query proteins. We select one representative structure for each of the six largest CATH architectures clusters, and randomly sample 10 latent codes based on the inferred posterior distribution with a temperature of 0.1. The sampled structures exhibit a level of similarity in global topology arrangement, as well as certain residue-level diversity.

This experiment also demonstrates another noticeable advantage of our model. One major reason for the non-uniform designability arises from the complex composition of the CATH-60 dataset [33] used for model training, which comprises a proportion of disordered structures. Although the encoder successfully learns to encode them into a specific region in the latent manifold, by default, the latent diffusion model and the generative diffusion model still collectively learn to faithfully sample around those highly disordered proteins. Since these samples and high-quality ones have distinct latent distributions, we could freely choose to include or exclude the sampling of those regions with the assistance of additional latent classifiers or guidances. Empirically, a cutoff of 0.6 would prohibit sampling these regions with only minor decreases on the other metrics. Hence, as long as the model has sufficient expressivity, the existence of noisy or bad samples will have a limited effect on the model’s learning of high-quality samples. That is, the model is likely to exhibit higher robustness to noisy training data.

### G Controlling topologies of sampled proteins with specified query proteins

